# Huntingtin lowering reduces somatic instability at CAG-expanded loci

**DOI:** 10.1101/2020.07.23.218347

**Authors:** Sydney R. Coffey, Marissa Andrew, Heather Ging, Joseph Hamilton, Michael Flower, Marina Kovalenko, Robert M. Bragg, Jeffrey P. Cantle, Cassandra A. McHugh, José M. Carrillo, Julie-Anne Rodier, Deanna M. Marchionini, Hilary A. Wilkinson, Seung Kwak, David S. Howland, C. Frank Bennett, Ricardo Mouro Pinto, Georg Auburger, Scott O. Zeitlin, Holly B. Kordasiewicz, Sarah J. Tabrizi, Vanessa C. Wheeler, Jeffrey B. Carroll

**Affiliations:** Behavioral Neuroscience Program, Psychology Department, Western Washington University, Bellingham, WA 98225, USA; Molecular Neurogenetics Unit, Center for Genomic Medicine, Massachusetts General Hospital, Boston, MA 02114, USA; UCL Huntington’s Disease Centre, Department of Neurodegenerative Disease, UCL Queen Square Institute of Neurology, Queen Square, London, WC1N 3BG, UK; UK Dementia Research Institute, University College London, WC1N 3BG, UK; CHDI Foundation/CHDI Management, NJ 08540, USA; Ionis Pharmaceuticals, Carlsbad, CA 92008, USA; Department of Neurology, Harvard Medical School, Boston, MA 02114, USA; Experimental Neurology, Medical School, Goethe University, 60590 Frankfurt am Main, Germany; Department of Neuroscience, University of Virginia School of Medicine, Charlottesville, VA 22903, USA

**Author notes:** **Corresponding Authors:** Vanessa Wheeler, Jeff Carroll.

## Abstract

Expanded trinucleotide repeats cause many human diseases, including Huntington’s disease (HD). Recent studies indicate that somatic instability of these repeats contributes to pathogenesis in several expansion disorders. We find that lowering huntingtin protein (HTT) levels reduces somatic instability of both the *Htt* and *Atxn2* CAG tracts in knockin mouse models, and the *HTT* CAG tract in human iPSC-derived neurons, revealing an unexpected role for HTT in regulating somatic instability.

## Main Text

Many expanded trinucleotide repeats are unstable in somatic cells ^1,2^. Instability is expansion-biased and occurs in a repeat length-, time- and cell-type-dependent manner. A role for somatic instability in HD pathology is strongly supported by genome wide association studies of HD patients examining the residual age of onset ^3,4^ or a measure of disease progression ^5^ after correcting for CAG length. These studies revealed that genetic variation in DNA repair genes, in particular mismatch repair (MMR) genes and the structure-specific nuclease FANCD2 and FANCI associated nuclease 1 (*FAN1*), altered the rate of onset or progression^3,5^. MMR genes are essential drivers of somatic CAG.CTG expansion ^6–8^ and although FAN1 is not a canonical MMR protein, its loss is associated with exacerbated repeat expansion in both *HTT* ^9^ and *FMR1* ^10^. Single nucleotide polymorphisms (SNPs) in some of these genes were also associated with residual age of onset in spinocerebellar ataxias (SCAs), supporting somatic expansion as driver of pathogenesis across multiple CAG expansion disorders. Additional key support for the importance of somatic expansion in HD pathogenesis comes from studies of rare *HTT* haplotypes with either the duplication or loss of the penultimate CAACAG sequence at the 3’ end of the CAG tract, demonstrating that pure CAG tract length, rather than polyglutamine length, drives the rate of HD onset ^4,11^. This supports observations from other repeat diseases, including myotonic dystrophy type 1 ^12^ and SCA types 1 ^13^ and 2 ^14^, where interruptions of the expanded CTG/CAG repeat modulate age of onset. Collectively, the emerging picture indicates that somatic instability, driven by DNA repair gene-mediated action on pure repeats, is a key driver of the rate of pathogenic process in multiple repeat expansion disorders.

Because clinical investigations of huntingtin HTT-lowering treatments are rapidly advancing ^15^ and somatic instability may contribute substantially to HD pathogenesis, we wondered if HTT-lowering therapies could influence somatic instability. We suppressed HTT in peripheral tissues of knock-in HD mouse models (*Htt*^*Q111/+*^) via repeated administration of antisense oligonucleotides (ASOs) targeting mouse *Htt* ^16^. Treatment with a pan-*Htt*-targeting ASO (hereafter *Htt* ASO) from 2 to 10 months of age reduces hepatic HTT levels by 64% ^16^ and is accompanied by a 37% reduction in the expansion index of the *Htt*^*Q111*^ CAG tract in the liver (Fig. 1A, middle panel). Somatic expansion in the striatum of these mice was unaffected by peripherally administered ASOs (Fig. S1, ANOVA treatment effect: F_(2,8)_ = 0.68, p = 0.53), which are excluded by the blood brain barrier ^17^. After our initial observation, we subsequently generated two additional cohorts of ASO-treated animals sacrificed after 5.5 or 12 months of treatment - timepoints flanking our initial cohort’s treatment duration - in which we replicated our finding that decreased HTT levels are associated with decreased somatic expansion in the liver (Fig. 1A: N = 57 animals, 3-9 per treatment arm over three ages, ANOVA for treatment effect F_(2,48)_ = 203.65, p < 2.2e-16; exemplar PCR product traces shown in Fig. 1B). This decreased somatic expansion was not attributable to inherited allele sizes, which were similar across treatment groups (Table S1, ANOVA treatment effect F_(2, 54)_ = 0.42 p = 0.66). Increased duration of HTT suppression was associated with greater reductions in somatic expansion, with expansion indices reduced by 50% in the liver after 12 months of treatment.

**Figure 1:**
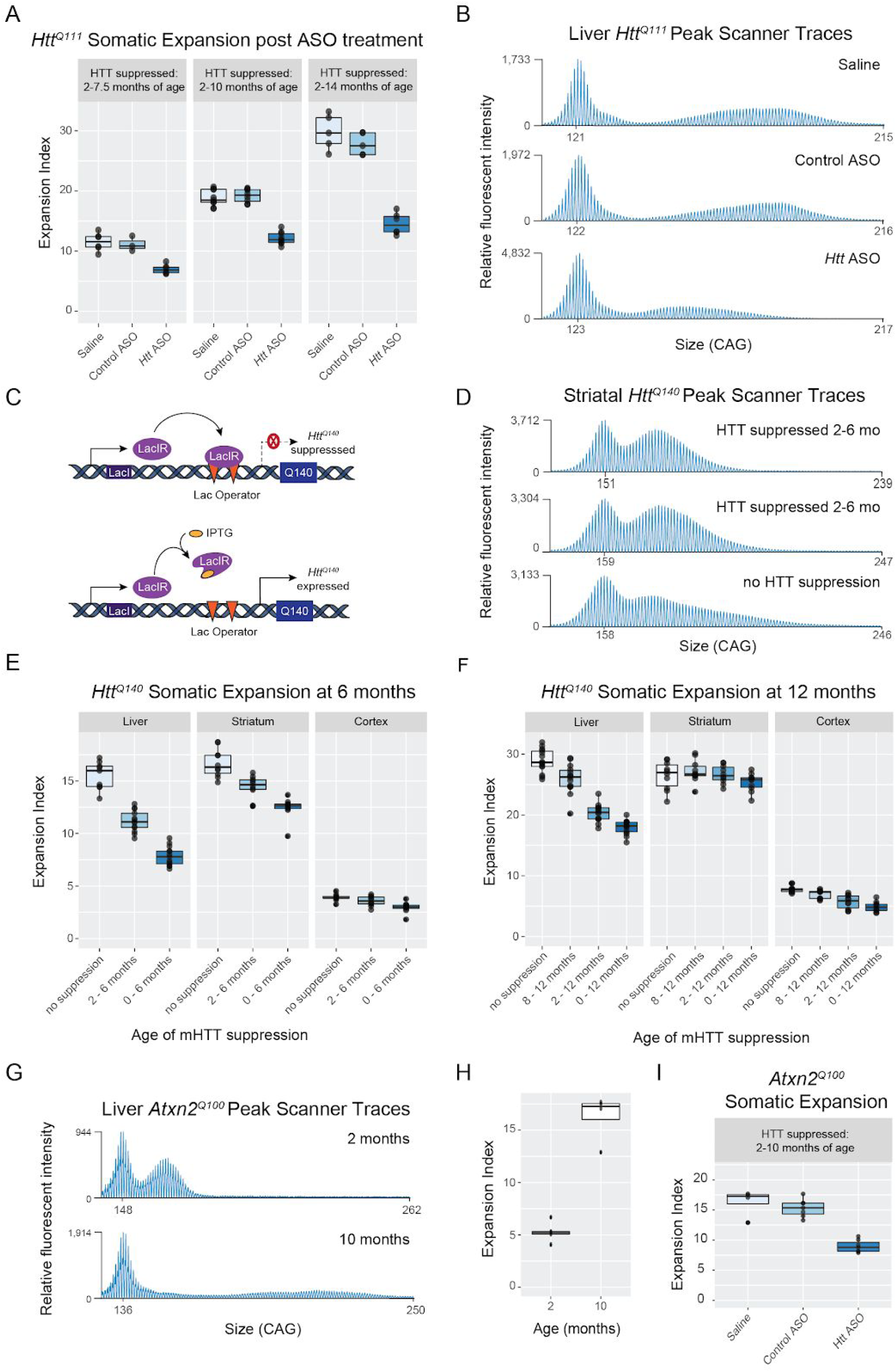
Huntingtin lowering reduces somatic expansion of expanded *Htt* and *Atxn2* CAG tracts. (A) Chronic peripheral treatment for the indicated intervals with *Htt* ASO reduces somatic expansion in the liver of *Htt*^*Q111/+*^ mice compared to control ASO or saline alone, neither of which had an effect on HTT levels or the expansion index (ANOVA treatment effect over a total of 57 mice: F_(2,48)_ = 203.65, p < 2.2e-16). (B) Exemplar traces of *Htt* CAG repeat-containing PCR products from *Htt*^*Q111/+*^ mouse livers highlights the reduction in the expanded repeat population in mice treated with *Htt* ASOs for 12 months. (C) Schematic of the regulatable *Htt*^*LacO-Q140*^ allele. The transcriptional repressor, LacIR, binds the Lac operator, preventing transcription of the *Htt*^*LacO-Q140*^ allele (top). In practice, we achieved approximately 50% reduced expression of the *Htt*^*LacO-Q140*^ allele with a concomitant 50% knockdown of mutant HTT. LacIR is allosterically inhibited by isopropyl β-d-1-thiogalactopyranoside (IPTG), enabling expression of the *Htt*^*LacO-Q140*^ allele (bottom). (D) Exemplar Peak Scanner 2 traces of *Htt* CAG repeat-containing PCR products from 6-month-old *Htt*^*LacO-Q140/+*^ striata illustrate the reduction in the expanded repeat population upon suppression of the *Htt*^*LacO-Q140*^ allele. (E,F) Approximately 50% mutant HTT reduction for the indicated durations, resulted in a time-dependent reduction in somatic expansion in 6-month old *Htt*^*LacO-Q140/+*^ mice (ANOVA treatment effect: F_(2,86)_ = 148.28, p < 2.2e-16.) (E) or in 12-month old *Htt*^*LacO-Q140/+*^ mice (F). (ANOVA treatment effect: F_(2,123)_ = 91.11, p < 2.2e-16). (G) Exemplar traces of *Atxn2* CAG repeat-containing PCR products from *Atxn2*^*Q100/+*^ mouse livers at 2 and 10 months of age. (H) Expansion indices for the *Atxn2*^*Q100/+*^ CAG repeat in 2- and 10-month old liver reveal an increase with age (Welch two sample t-test: t_(11.67)_ = 4.474, p = 5.15e-4). (I) Treatment with *Htt* ASO reduces somatic expansion in *Atxn2*^*Q100/+*^ livers (ANOVA treatment effect: F_(2,15)_ = 65.54, p = 2.59e-08). All exemplar Peak Scanner 2 traces are displayed spanning equal CAG distances within each figure and aligned to the modal allele.

To determine whether this effect was occurring in an orthogonal genetic system, we quantified somatic expansion in mice in which expression levels of another knockin CAG-expanded *Htt* allele (Q140)^18^ are regulatable via an upstream LacO repressor binding site, the *Htt*^*LacO-Q140/+*^ mice (Fig. 1C, Marchionini *et al*., in preparation). This renders the *Htt*^*Q140*^ allele partially repressible by expression of a *lacI* repressor, driven by the broadly expressed β-actin promoter ^19^. Treatment with isopropyl β-d-1-thiogalactopyranoside (IPTG) recruits the lac repressor away from the targeted locus, enabling expression of the targeted gene. In practice, we observe approximately 50% reduction in mutant HTT expression throughout the body upon IPTG withdrawal (Marchionini *et al*., in preparation). In our initial *Htt*^*Q111/+*^ cohorts, we began ASO treatment at 2 months of age - before the emergence of most disease-like phenotypes ^20^ and when low levels of somatic expansion are present ^21^. To investigate whether HTT-lowering interventions initiated at a later stage of disease influence somatic instability, we tested a range of mHTT suppression durations, starting at either 0, 2 or 8 months of age. Within each cohort, prolonging mHTT suppression led to greater reductions in somatic expansion of the targeted *Htt*^*Q140*^ allele (Fig. 1E-F, ANOVA treatment effect at 6 months of age F_(2,86)_ = 148.28, p < 2.2e-16; and at 12 months of age F_(2,123)_ = 91.11, p < 2.2e-16; exemplar striatal PCR product traces in 1D). The reduced expansion indices did not reflect differences in inherited allele sizes, which were similar across groups (Table S2). This effect is seen both in the liver and striatum, confirming that the relationship between somatic instability and HTT levels is not limited to peripheral organs or to ASO treatment.

To determine whether HTT plays a general role in somatic instability, or whether this role is limited to its own locus, we turned to a recently described knockin model of of spinocerebellar ataxia type 2 (SCA2) containing an expanded CAG repeat in the *Atxn2* locus (*Atxn2*^Q100/+^)^22^. These mice have progressive molecular and physiological phenotypes, and between 2 and 10 months of age we observe robust, somatic expansion of the *Atxn2*^*Q100*^ CAG tract in the liver (Fig. 1G-H, Welch two sample t-test: t_(11.67)_ = 4.5, p = 5.15e-4). To mirror our initial cohort of ASO-treated *Htt*^*Q111/+*^ mice, we injected *Atxn2*^*Q100/+*^ mice from 2-10 months of age with the same *Htt* ASO used in our *Htt* studies. At sacrifice, we observe robust reductions in hepatic HTT in *Htt* ASO-treated *Atxn2*^Q100/+^ mice (Fig S2A-B, ANOVA for treatment effect: F_(2, 9)_ = 13.8, p=1.8e-3) with no change in ataxin-2 protein or transcript levels (Fig S2C-E, ANOVA for treatment effect on protein: F_(2, 21)_ = 0.47, p = 0.63; and transcripts: F_(2,10)_ = 1.81, p = 0.21). Similar to the impact on the *Htt*^*Q111*^ CAG repeat, the expansion index of the *Atxn2*^Q100^ CAG repeat was reduced by 43% in *Htt* ASO-treated mice (Fig. 1I, ANOVA treatment effect: F_(2,15)_ = 64.5, p = 4.3e-08). Reduced somatic expansion in the *Htt* ASO-treated mice was not attributable to differences in inherited allele size, which was similar across treatment groups (Table S1, ANOVA treatment effect: F_(2,15)_ = 1.69, p =0.22). These results demonstrate that HTT-lowering-mediated suppression of somatic expansion can occur *in trans*, and in the absence of changes in the level of the gene product of the unstable locus. Further, the comparable impacts of *Htt* ASO treatment on the *Htt*^*Q111*^ and *Atxn2*^*Q100*^ CAG repeats demonstrate that somatic expansion is promoted by a function of wild-type HTT rather than by a toxic property specific to mutant HTT.

To examine the effect of ASO-mediated HTT knockdown in human medium spiny neurons (MSNs), the cell type most vulnerable in HD ^23^, we generated induced pluripotent stem cells (iPSC) from a juvenile-onset HD patient and differentiated them into MSN-containing cultures with approximately 137 CAG repeats (Fig. 2A) ^24^. MSN fate in untreated, parallel cultures was confirmed after 112 days in culture by high-content imaging. Cultures contained 53% neurons (SD ± 11.34), of which 48% (SD ± 18.96) were positive for DARPP-32 (Fig. 2B, n = 5 technical replicates). We treated mature, MSN-containing cultures on culture day 37 (treatment day 0) with 40 μM *HTT*-targeting ASO or vehicle for 98 days (Fig. 2A). In parallel cultures we confirmed consistent *HTT* transcript knockdown of greater than 70% over 35 days of *HTT* ASO treatment (Fig. S3, ANOVA treatment effect: F_(15,32)_ = 16.24, p = <0.0001). We calculated the modal allele shift, the instability index ^25^, and a composite index that captures changes in both the modal allele and the repeat length distribution over time (Fig. 2C). Unlike the instability index, which measures CAG distance relative to the modal allele within each sample, the composite index measures CAG distance relative to the modal allele of each biological replicate early in treatment (day 3). In this way, the composite index potentially provides a more sensitive endpoint to test modifiers that might impact different aspects of somatic instability. We found that the composite index was reduced in cultures receiving *HTT* ASO compared to vehicle control (Fig. 2D, N=2 independent differentiations over 6 timepoints, linear mixed effects model treatment effect: F_(1,19)_ = 29.31; p = 3.18e-5). The reduced composite index could not be explained by shifts in the instability index or modal allele alone (Fig. 2E-F, linear mixed effects model treatment effect on the instability index: F_(1,19)_ =1.1; p = 0.31; and modal allele shift: F_(1,19)_ = 1.5; p = 0.24). These results indicate that HTT is involved in expansion of its own CAG repeat in both mouse models and human iPSC-derived MSNs.

**Figure 2:**
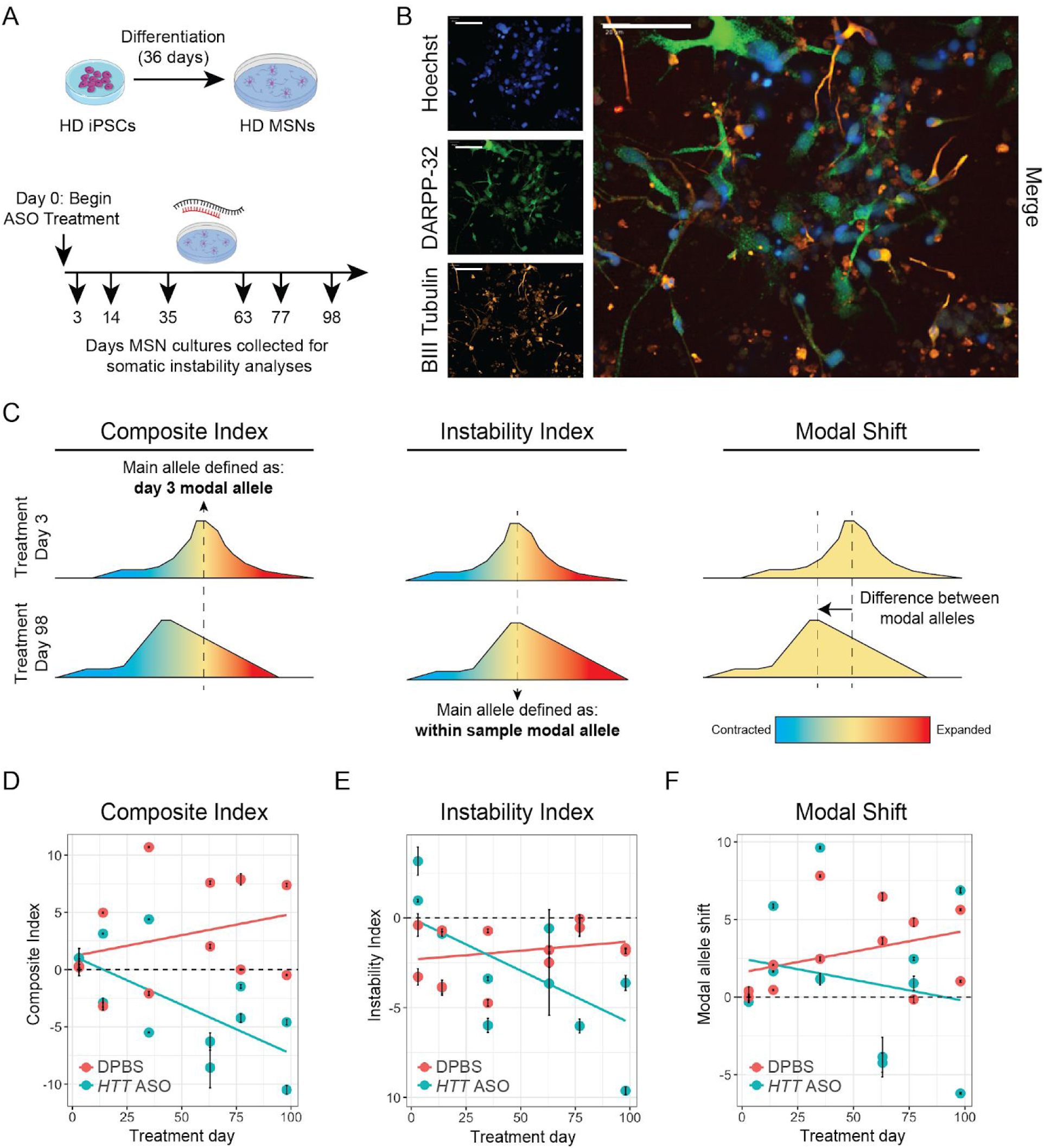
ASO-mediated HTT knockdown reduces somatic instability of the *HTT* CAG tract in human MSN-containing cultures. (A) Schematic of 36 day MSN differentiation protocol and subsequent 40 µM *HTT* ASO treatment beginning at culture day 37 (treatment day 0) and continuing for 98 days. MSN-containing cultures were collected at 3, 14, 35, 63, 77, and 98 days into treatment for somatic instability analyses. (B) In untreated, parallel cultures high content imaging performed at culture day 112 determined that of the β-III Tubulin positive viable cells, 47.82% co-stained for DARPP-32 (n = 5 technical replicates). DARPP-32, β-III Tubulin, and Hoechst are markers of striatal MSNs, neurons, and nuclei, respectively. Scale bars represent 60 µm. (C) Graphical representation of the three different measures calculated. Both the composite and instability indices capture the distribution of repeat lengths by calculating the CAG distance from the main allele. In the composite index (left), the main allele is defined as the treatment day 3 modal allele within each MSN differentiation, whereas for the instability index (middle), the main allele is defined as the within-sample modal allele. In this way, the composite index encompasses changes in both the modal allele and the distribution of CAG tract lengths, while the instability index only captures changes in the allele distribution relative to the modal allele. Modal allele shifts (right) were calculated relative to treatment day 3. (D) There was a significant difference between the composite index of *HTT* ASO-treated HD MSN-containing cultures compared to DPBS-treated cultures (ANOVA treatment effect: F_(1,19)_ = 29.31; p = 3.18e-5). Data from each time point in D-F represent two biological replicates (independent MSN differentiations) with an average of 8 technical replicates. Linear models have been fitted, with *HTT* ASO treatment in red and DPBS treatment in teal. Error bars represent standard deviation (SD).

Our results suggest that HTT-lowering treatments, which are rapidly progressing through clinical trials ^15^, may have the unexpected benefit of reducing somatic instability of the *HTT* CAG repeat. They reduce instability in at least one other repeat expanded locus (*Atxn2*), suggesting they are worth considering as treatments in other repeat expansion diseases - a large family of currently untreatable conditions ^26^. Many disease-associated repeats exhibit somatic expansion ^27^ that is thought to play a role in driving the rate of pathogenesis ^28^. Therefore, to the extent that repeat expansion in relevant cell types in these disorders is sensitive to HTT levels, HTT-lowering treatments may prove effective in multiple indications. Our data are consistent with multiple, non-exclusive mechanisms by which HTT could influence somatic instability of CAG repeats. HTT has been proposed to scaffold the enzymes involved in transcription-coupled DNA repair complexes and the critical DNA repair kinase ATM ^29^, which repairs DNA after oxidative damage ^30^.^30^. HTT also plays widely observed, but poorly delineated, roles in regulating transcription and chromatin states, including post-translational modifications of histones ^31,32^. It has been suggested that transcription is required for repeat instability ^33^ though this remains untested in animal models. As we do not observe reduced *Atxn2* transcripts in the *Htt* ASO treated mice, it appears unlikely that HTT promotes repeat instability by impacting transcription at expanded repeat loci themselves. Alternatively, HTT may regulate chromatin states of these loci, influencing access of DNA repair proteins controlling repeat instability or influencing expression levels of DNA repair proteins themselves ^33^. While the relative contributions of these mechanisms remain to be determined, our observations clearly demonstrate that HTT positively regulates somatic CAG expansion at *Htt/HTT* and *Atxn2* loci, and that this mechanism may provide cross-disease utility in the treatment of repeat expansion diseases.

## Methods

### Mice

All mice in the ASO treatment cohorts were group housed in the Western Washington University (WWU) vivarium with *ad libitum* access to food and water. *Htt*^*Q111/+*^ mice (RRID:IMSR_JAX:003456) in the 5.5-month treatment cohort were bred at WWU and *Htt*^*Q111/+*^ mice in the 8-month and 12-month treatment cohorts were acquired from Jackson Laboratories (Bar Harbor, ME) at 1.5 months of age. *Atxn2*^*Q100/+*^ breeders were provided by Georg Auburger’s lab at Goethe University Medical School Frankfurt with subsequent experimental mice bred at WWU. Inherited allele sizes of all experimental mice were determined from tail tissue taken at weaning (21 days) at Laragen, Inc (Table S1). Experimental N for each of the *Htt*^*Q111/+*^ and *Atxn2*^*Q100/+*^ cohorts are reported in Table S1. Beginning at 63 (±11, SD = 3.6) days of age, female *Htt*^*Q111/+*^ and *Atxn2*^*Q100/+*^ mice were intraperitoneally (IP) injected with 50 mg per kg body weight (mpk) of pan *Htt* targeted ASO (‘*Htt* ASO’), a control ASO without a target in the mouse genome (‘Control ASO’), or 4 µL/g body weight of saline alone. ASOs were synthesized by Ionis Pharmaceuticals (Carlsbad, CA). Sequence information and chemistry of both the *Htt* and control ASOs were published previously ^16^. Cohorts of *Htt*^*Q111/+*^ mice (data presented in Fig. 1A-B) were treated weekly for 5.5 (169 ± 2 days), 8 (237 ± 4 days), or 12 (366 ± 4 days) months. Cohorts of *Atxn2*^Q100/+^ mice (data presented in Fig. 1G-I) were treated weekly for 8 (261 ± 7 days) months. *Atxn2*^*Q100/+*^ and *Htt*^*Q111/+*^ mice in the 5.5- and 12-month treatment cohorts were IP injected with 180 mpk 5-ethynyl uridine (Click Chemistry Tools) 5 hours prior to sacrifice via lethal IP injection of at least 250 mpk sodium pentobarbital containing euthanasia solution. Liver, kidney, tail, striatum, cortex, and cerebellum were harvested and flash frozen in liquid nitrogen. All procedures performed at WWU were approved by the institutional animal care and use committee (protocol 16-011).

*Htt*^*LacO-Q140/+*^; *b-actin-LacI*^*R*^ *tg* mice (hereafter *Htt*^*LacO-Q140/+*^ mice) were housed at Psychogenics with *ad libitum* access to food and water. For maintaining normal levels of mutant HTT expression, IPTG (2.4 mg/mL) was added to the drinking water of dams pregnant with *Htt*^*LacO-Q140/+*^ pups starting at embryonic day 5. At weaning *Htt*^*LacO-Q140/+*^ mice were treated continuously with IPTG (2.4 mg/mL) in their drinking water for the duration of the experiment (6 or 12 months of age) or until either 2 or 8 months of age (Figures 1 E & F). Maximal mutant HTT protein suppression occurred approximately 10 days following IPTG withdrawal. Approximately equal numbers of male and female mice were used to quantify somatic expansion (See Table S2 for detailed cohort characteristics). Genotyping and inherited allele sizing was performed at Laragen, Inc. using weaning tail tissue. All procedures performed at Psychogenics were approved by the institutional animal care and use committee (protocol 271-0315).

### iPSC Cultures

125 CAG iPSCs were generated from peripheral blood mononuclear cells (PBMCs) using the Sendai CytoTune™-iPS 2.0 method at the National Hospital for Neurology and Neurosurgery (Queen Square, London) and Censo Biotechnologies. Illumina targeted sequencing determined that this individual has canonical *HTT* CAG alleles, with one terminal CAACAG interruption. Stem cells were maintained in Essential 8™ media (Thermo) on Nunc™ plasticware (Thermo), coated with Geltrex™ (Gibco) diluted 1:100 with DMEM:F-12. 125 CAG iPSCs were passaged weekly when they reached 80% confluency with 0.02% EDTA (Gibco). Because repeat expansion occurs during maintenance of iPSC lines, we re-sized mature MSNs at treatment day 3, finding an average CAG tract length of 137. Differentiation into MSN-containing cultures was achieved using activin A to direct striatal fate as previously described ^24^. Briefly, iPSCs were guided towards a neuroectodermal fate by maintaining cells in Pre-26 media (1/3 Neurobasal, 2/3 DMEM:F-12, 1:100 L-glutamine, 1:150 N2, 1:150 B27 without Vitamin A, 100μM β-mercaptoethanol) with triple SMAD inhibition (200mM dorsomorphin, 100mM LDN193189, 10μM SB431542) for 9 days. Cells were passaged with 0.02% EDTA onto fibronectin (1:40 in DPBS) and maintained in Pre-26 media containing 25 ng/mL activin A to induce a striatal/ lateral ganglionic eminence (LGE) fate. After 10 days, cells were passaged onto laminin (1:50 in DMEM:F-12) and cultured in Pre-26 media plus activin A for a further 7 days. At this point, cells were terminally maintained in Post-26 media (1/3 Neurobasal, 2/3 DMEM:F-12, 1:100 L-glutamine, 1:150 N2, 1:150 B27 with Vitamin A, 100μM β-mercaptoethanol) supplemented with 25 ng/mL activin A, 10 ng/mL BDNF and 10 ng/mL GDNF. At resting state, these MSNs are unstable over 100 days in culture with a change in modal CAG repeat length of +0.026 CAG per day and an increase in the composite index of 0.033 CAG per day.

Ionis Pharmaceuticals (Carlsbad, CA) provided the *HTT* ASO and scrambled ASO. 200 mg powder ASOs were reconstituted in 1 mL DPBS and filtered through a 0.22 µM syringe filter. Stock concentration was determined by measuring absorbance at 260 nm, using the formula ((260 nm absorbance x dilution factor x molecular weight) / (extinction coefficient x 1000)). For somatic instability experiments, mature HD MSNs were treated with 40 µM *HTT* ASO or DPBS starting on culture day 37. To assess *HTT* knockdown, parallel HD MSN cultures were treated beginning at culture day 37 with 40 µM *HTT* ASO, scrambled ASO, or DPBS. *HTT* ASO, scrambled ASO, or DPBS was delivered in Post-26 media during twice-weekly media changes. Samples were collected at 3, 14, 35, 63, 77 and 98 days into treatment for somatic instability analyses. For qRT-PCR, cells were collected at 0, 3, 7, 14, 28, and 35 days into treatment. Untreated MSN cultures were collected at culture day 112 for high content imaging. At each timepoint, cells were lifted using 0.5 mL Accutase (Stemcell Technologies), then centrifuged at 1,000 rpm and Accutase removed. They were washed in 0.5 mL DPBS, then the cell pellet was stored at −80°C. gDNA was extracted using the Nanobind kit (Circulomics).

### PCR and fragment analyses

Genomic DNA from *Htt*^*Q111/+*^ and *Atxn2*^*Q100/+*^ mice was extracted from 10-30 mg of liver tissue via phenol-chloroform extraction. Liver tissue was incubated overnight at 55°C in 0.5 mL of lipid rich tissue buffer (50 mM Tris, 100 mM EDTA, 2% SDS, pH 8.0) plus 35 µL of 20 mg/mL proteinase K (Thermo Fisher Scientific cat# AM2544). Liver tissue was homogenized by pestle, then 500 µL of phenol:chloroform:isoamyl-alcohol (25:24:1; Thermo Fisher Scientific cat# 15593-049) was added to the liver lysate. Samples were shaken for 5 minutes then centrifuged at 16,000 g for 5 minutes. Following centrifugation, the supernatant was transferred to a clean tube, and 50 µL of 3 M sodium acetate was added to the lysate. Then 850 µL of absolute ethanol was added to the lysate and gently mixed to precipitate the DNA. Samples were centrifuged for 15 minutes at 16,000 g, then the supernatant was decanted. Pelleted DNA was washed with 1 mL of 70% ethanol and centrifuged for 15 minutes at 16,000 g. The supernatant was removed, and residual ethanol was allowed to evaporate for 1 hour. Pelleted DNA was re-suspended by incubating in 100 µL of water at room temperature overnight or at 55°C for 1 hour. Genomic DNA concentrations were determined using the Synergy 2 Plate Reader (BioTek). Genomic DNA from the striatum, cortex, and liver of *Htt*^*LacO-Q140/+*^ mice was extracted and DNA concentrations determined by Q^2^ Solutions.

*Htt* CAG repeat-containing PCR products from both the *Htt*^*Q111/+*^ and *Htt*^*LacO-Q140*^ cohorts were amplified in 80 µL PCR reactions containing 8 µL DMSO, 4 µL 10 mg/mL BSA, 8 µL 2 mM dNTPs (Thermo Fisher Scientific cat# R0241), 6.4 µL 10 µM 5’ 6-FAM-labelled CAG1 primer ^34^, 6.4 µL 10 µM HU3 primer ^34^, and 200 ng genomic DNA, in addition to the following components of the AmpliTaq 360 DNA Polymerase kit (Fisher Scientific, cat no. 43-988-18): 8 µL AmpliTaq Buffer, 8 µL GC Enhancer, 3.2 µL 25 mM MgCl_2,_ and 0.8 µL AmpliTaq Taq. Thermocycling conditions were as follows: an initial denature at 94°C for 90 s, 35 repeated cycles of 94°C for 30 s, 63.5°C for 30 s, and 72°C for 90 s, with a final extension at 72°C for 10 minutes.

*Atxn2* PCR reactions consisted of 6.4 µL 5 µM 5’ 6FAM-labelled SCA2Ex1-Fwd5 primer ^22^, 6.4 µL 5 µM SCA2Ex1-Rev2 ^22^, 8 µL 25 ng/µL genomic DNA, 31.6 µL water and the following reagents from the Taq PCR Core Kit (Qiagen, cat no. 201223): 1.6 µL 25 mM MgCl_2_, 8 µL 10x buffer, 16 µL 5x Q Reagent, 1.6 µL dNTPs (10 mM each), and 0.4 uL Taq. A hot-start PCR, where the thermocycler was brought to 95°C before thermocycling began, was performed. After an initial denature at 95°C for 5 minutes, a 9-cycle touchdown phase began, consisting of a 30 s denature step at 95°C, 30 s annealing step starting at 70°C and decreasing by 1°C each cycle, and a 90 second extension step at 72°C. After the touchdown phase, traditional thermocycling continued for 32 cycles of 95°C for 30 s, 62°C for 30 s, and 72°C for 90 s. A final extension at 72°C for 10 minutes concluded thermocycling.

*Htt* and *Atxn2* amplicons generated from *Htt*^*Q111/+*^, *Htt*^*LacO-Q140/+*^, and *Atxn2*^*Q100/+*^ cohorts were concentrated from 80 µL to 20 µL using the GeneJET Purification Kit (Thermo Fisher Scientific, cat no. K0702). Two minimal modifications to the manufacturer’s protocol were made, specifically PCR products were eluted in 20 µL of water, instead of the elution buffer, and the flow-through was run through the column twice. Concentrated PCR products were sent to Genewiz for fragment analysis.

CAG repeat-spanning *HTT* PCR products were amplified from HD MSN genomic DNA in 100 µL reactions containing 50 µL AmpliTaq Gold 360 Master Mix, 10 µL GC enhancer, 4 µL each of 5 µM stock forward primer (HD3F: CCTTCGAGTCCCTCAAGTCCTT) and reverse primer (HD5: CGGCTGAGGCAGCAGCGGCTGT), 1 µL 50 ng/µL gDNA, and 28 µL dH_2_O with the following cycling conditions: 94°C for 90 s, 35 cycles of 94°C for 30 s, 65°C for 30 s and 72°C for 90 s, and the final extension at 72°C for 10 mins. PCR products were cleaned and concentrated with AMPure XP beads prior to capillary electrophoresis. Samples (2 µL) were suspended in 10 µL Hi-Di formamide (Thermo Fisher) and 0.5 µL MapMarker ROX 1000 (Eurogentec) size standard on a low profile non-skirted 96 well plate and heated to 95°C for 4 min, then cooled to 4°C before being resolved by capillary electrophoresis on an ABI 3730xl Genetic Analyser.

### Somatic instability analyses

PCR product sizes from the *Htt*^*Q111/+*^, *Htt*^*LacO-Q140/+*^, and *Atxn2*^*Q100/+*^ cohorts were assigned using the open-source software Peak Scanner 2 or GeneMapper v5 (both produced by Thermo Fisher Scientific). Expansion indices were calculated following previously described methods ^25^, but in our analyses we defined the main allele as the modal allele on a per sample basis, rather than the peak with the greatest fluorescence in tail PCR products from matched mice. Under this definition, CAG distances were determined based on the distance from the modal allele within each sample and expanded alleles were considered peaks with positive CAG distances. Samples in which the fluorescence of the modal allele was below 800 relative fluorescent units (RFU, arbitrary units) were considered to have failed quality controls and were omitted from analyses. A 10% threshold was applied to the *Htt*^*Q111/+*^ and *Htt*^*LacO-Q140/+*^ samples and a 5% threshold was applied to the *Atxn2*^*Q100/+*^ samples. To account for the faster migration speed of CAG repeats compared to random sequences in capillary electrophoresis ^35^, CAG sizes reported in exemplar traces (Fig 1B and G) were calculated by (1) subtracting the base pair length of the regions flanking the CAG repeat from the overall amplicon size as determined by Peak Scanner 2, (2) converting to CAG units by dividing the corrected bp length by 3, (3) multiplying the resulting CAG length by 1.045, and (4) adding 1.2088 CAGs. Capillary electrophoresis chromatographs from the HD MSN-containing cultures were aligned in GeneMapper v5 software. To measure modal CAG repeat length and instability index ^25^, data was exported and analysed with a custom R script, available at https://caginstability.ml, using an inclusion threshold of 10% of modal peak height, and confirmed manually. Linear regression models were built in R and compared by an analysis of covariance (ANCOVA).

### Western blotting

Liver protein was extracted from 40-55 mg of tissue in 500 μL of RIPA buffer (25 mM Tris-HCl, 150 mM NaCl, 1% NP-40, 1% sodium deoxycholate, 0.1% SDS) with protease and phosphatase inhibitors added (Thermo Fisher Scientific, cat no. 78442). Liver tissue was placed in BeadBug tubes loaded to ⅓ capacity with 1.4 mm Ceramic Beads (Omni-International cat no. 19-645) and homogenized using the Bead Blaster 24 (Benchmark) for 2 cycles at 7m/s with cycles consisting of 2 × 30 seconds with a 10 second intermission. Following homogenization, samples were centrifuged at 20,000 g for 15 minutes at 4°C, and the supernatant was collected. Protein concentration was determined by Pierce BCA assay (Thermo Fisher Scientific, cat no. 23225) according to the manufacturer’s protocol. Prior to western blotting, 100 μg of liver protein were mixed with 4x loading buffer (ThermoScientific, cat no. NP007) and 10x reducing agent (Thermo Fisher Scientific, cat no. NP004), then denatured for 10 minutes at 70°C. Protein lysates were loaded into 3-8% Tris-Acetate mini gels (Thermo Fisher Scientific, cat no. EA03785BOX), separated electrophoretically, and transferred to a PVDF membrane (Millipore, cat no. IPFL00010) for 17-20 hours at 4°C. Membranes were stained using the Revert 700 Total Protein Stain Kit (Li-Cor, cat no. 926-11016) following the manufacturer’s protocol, then imaged with the Li-Cor Odyssey Fc Instrument to enable normalization to total protein signal. After destaining, the membranes were blocked in Intercept TBS Blocking Buffer (Li-Cor, cat no. 927-70001) for 1 hour. Membranes were incubated overnight at 4°C in primary antibodies targeting either ATXN2 or HTT. Primary antibodies used were polyclonal, rabbit anti-ATXN2 antibodies (Proteintech, cat no. #21776-1-AP) and monoclonal, rabbit anti-huntingtin antibody EPR5526 (Abcam ab109115) diluted 1:500 and 1:1,000 respectively in Intercept blocking buffer plus 0.05% tween. Membranes were rinsed three times with TBS plus 0.05% tween before incubating for 1 hour at room temperature in 1:15,000 goat anti-rabbit (Li-Cor, cat no. 926-32211) secondary antibody diluted in Intercept blocking buffer, 0.05% tween, and 0.01% SDS. After rinsing three times with TBS plus 0.05% tween, membranes were imaged using the Li-Cor Odyssey Fc Instrument. Signal intensity was quantified in the ImageStudio Lite software, normalized to the total protein signal of the corresponding lane, and expressed relative to the average signal of saline-treated mice within the same blot.

### qRT-PCR

Liver tissue (30-50 mg) from the *Atxn2*^*Q100/+*^ mice was homogenized in 1 mL of Qiazol lysis reagent using the Bead Blaster 24 (Benchmark). Two cycles at 7 m/s were run, in which tissue was disrupted twice for 30 seconds, with a 10 second intermission. RNA was extracted using the RNeasy Lipid Tissue Mini kit (Qiagen 74804) according to the manufacturer’s protocol. cDNA synthesis and qRT-PCR was conducted as described previously ^20^, using a primer and probe set for *Atxn2* (Thermo Fisher, mM00485946_m1) that detects both the *Atxn2*^*Q1*^ and *Atxn2*^*Q100*^ alleles. RNA was extracted from the HD MSN-containing cultures using the RNeasy Plus Mini Kit (Qiagen). RNA concentration and quality was quantified on the NanoDrop Spectrometer and 0.5 to 2 μg of RNA was synthesised to cDNA using the SuperScript IV First-Strand Synthesis System (Thermo Fisher Scientific), as per manufacturer’s instructions. qRT-PCR was carried out with the TaqMan Fast Advanced Master Mix (Thermo Fisher Scientific) with the following reaction components; 7.5 μL master mix, 3 μL dH_2_O, 3 μL cDNA, and 1.5 μL probes. The reaction was combined in a 96-well plate and quantified on the QuantStudio 5 ProFlex system with the following thermal cycling conditions: 95°C for 40s, then 40 cycles of 95°C for 15s and 60°C for 30s. Data were analysed using the comparative Ct (cycle threshold) method (ΔΔCt). The geometric mean (geomean) of reference genes, *EIF4A2* (Hs00756996_g1), *UBC* (Hs00824723_m1), and *ATP5B* (Hs00969569_m1), was calculated and used to determine the relative expression of *HTT* (Hs00918174_m1). This was expressed relative to the average expression in the control samples. Fold change was given by 2^ ΔΔCt.

### High content imaging

Day 104 125Q MSNs were passaged into a 96 well format at 20K cells/well. The media was aspirated and the cells were dissociated with 0.5 ml Accutase per 1 well of a 12 well plate. The cell suspension was transferred to a 15 ml falcon tube containing PBS and a P1000 was used to triturate (6-10 passes) until a homogenous suspension was obtained. The cells were centrifuged at 300 xg for 3 min, then the PBS was aspirated and the pellet resuspended in 1 ml Post-26 media, with 1 µL Rock inhibitor (1:1000). The number of cells/ml were counted on the Tecan cell counter. A volume of cell suspension in fresh Post-26 media was prepared in a 50 ml Falcon tube sufficient to seed all plates required. 200 µL of 20K cells were plated per well in a 96 well plate that had been coated with Geltrex and incubated at 37°C for a minimum of 30 min. The cells were left untouched for 20 min before transferring to the incubator to allow the cells to settle and distribute evenly. After 48 h and twice per week thereafter, the cells underwent a half media change with 100 µL Post-26 media. The media was ‘spiked’ with Geltrex (1:50) for the first half media change to enhance cell attachment. Cells were fixed 1 week after passaging with 10% formalin solution containing 4% paraformaldehyde (Sigma) for 15 min at RT. The cells were washed 3 times with PBS and permeabilised with 0.2% Triton X-100 (Sigma) for 15 min at RT. Blocking solution containing 10% goat serum and 10% BSA in PBS was applied to the fixed cells for 1 h at RT. Primary antibodies (anti-DARPP-32 (rabbit, Abcam EP721Y ab40802, 1:200), anti-CTIP2 (rat, Abcam 25B6 ab18465, 1:300) anti-βIII tubulin (mouse 2G10 ab78078, 1:500) were added and incubated for 2 h in the dark at RT. Following four PBS washes, 100 µL of the secondary antibody, Alexa Fluor goat antibody (1:1000) (Invitrogen), was added for 1 h at RT in the dark. Following an additional PBS wash that was left on to incubate for 5 min at RT, 2X Hoechst solution (2 µL Hoechst 10mg/ml stock in 10 ml PBS) was added for 5 min at RT in the dark. A further three PBS washes were performed before adding 100 μl 0.02% sodium azide in PBS to the stained cells. HCI was performed using the Opera Phenix High Content Screening System, with 20 randomised fields-of-view captured per well. Images were analysed using Columbus image analysis software.

## Acknowledgments

We would like to thank Kimberly Cox and Washington Arias from PsychoGenics for their work on the *Htt*^*LacO-Q140/+*^ mouse studies.

## Declaration of Interests

### Research Funding

JBC has received research support from CHDI Foundation, Ionis Pharmaceuticals, Wave Life Sciences, and Triplet Therapeutics. Besides the authors of this paper who are employed by Ionis Pharmaceuticals (CFB, HK), no staff of these companies had any role in the design or interpretations of these experiments. SJT, HG and JH were funded by CHDI Foundation and DRI UK. MF was funded by DRI UK and Health Education England (HEE). JBC was funded by CHDI foundation and the Huntington Society of Canada. GA was funded by the DFG. RMP was the recipient of the Berman/Topper Family HD Career Development Fellowship from the Huntington’s Disease Society of America. SOZ was funded by the CHDI Foundation and by NIH NS090914. VCW was supported by the National Institutes of Health [NS049206] and by the CHDI Foundation.

### Employment

CFB and HBK are employees of Ionis Pharmaceuticals. JBC is a member of the scientific advisory board of Triplet Therapeutics.

### Personal financial interest

JBC is a member of the scientific advisory board of Triplet Therapeutics and has received consulting fees from Skyhawk Therapeutics. In the past 2 years, SJT has undertaken consultancy services, including membership of scientific advisory boards, with Alnylam Pharmaceuticals Inc., Annexon Inc., LoQus therapeutics (DDF Discovery Ltd), F. Hoffmann-La Roche Ltd, Genentech, PTC Bio, Novartis Pharma, Takeda Pharmaceuticals Ltd, Triplet Therapeutics, UCB Pharma S.A., University College Irvine and Vertex Pharmaceuticals Incorporated. All honoraria for these consultancies were paid through the offices of UCL Consultants Ltd., a wholly owned subsidiary of University College London. None of these companies had any role in the design or interpretations of these experiments. Other authors report no conflicts. VCW is a scientific advisory board member of Triplet Therapeutics, a company developing new therapeutic approaches to address triplet repeat disorders such Huntington’s disease and Myotonic Dystrophy, and of LoQus23 Therapeutics and has provided paid consulting services to Alnylam. Her financial interests in Triplet Therapeutics were reviewed and are managed by Massachusetts General Hospital and Partners HealthCare in accordance with their conflict of interest policies.

