## Supplementary Tables 1-2 for "Huntingtin lowering reduces somatic instability at CAG-expanded loci"

| Genotype | Treatment duration (months) | Sacrifice age (months) | Treatment | Sex | N | Average Inherited Allele Size (CAG no.) | Inherited Allele Size Standard Deviation |
|---|---|---|---|---|---|---|---|
| *Htt*$^{Q111/+}$ | 5.5 | 7.5 | *Htt* ASO | female | 6 | 117.8 | 5.4 |
| *Htt*$^{Q111/+}$ | 5.5 | 7.5 | Control ASO | female | 3 | 120.0 | 4.4 |
| *Htt*$^{Q111/+}$ | 5.5 | 7.5 | Saline | female | 6 | 120.2 | 7.4 |
| *Htt*$^{Q111/+}$ | 8 | 10 | *Htt* ASO | female | 9 | 114.3 | 3.4 |
| *Htt*$^{Q111/+}$ | 8 | 10 | Control ASO | female | 8 | 114.5 | 3.2 |
| *Htt*$^{Q111/+}$ | 8 | 10 | Saline | female | 9 | 114.9 | 2.8 |
| *Htt*$^{Q111/+}$ | 12 | 14 | *Htt* ASO | female | 6 | 117.7 | 0.8 |
| *Htt*$^{Q111/+}$ | 12 | 14 | Control ASO | female | 5 | 119.0 | 1.4 |
| *Htt*$^{Q111/+}$ | 12 | 14 | Saline | female | 5 | 119.0 | 1.0 |
| *Atxn2*$^{Q100/+}$ | 8 | 10 | *Htt* ASO | female | 7 | 135.9 | 2.3 |
| *Atxn2*$^{Q100/+}$ | 8 | 10 | Control ASO | female | 7 | 137.3 | 4.6 |
| *Atxn2*$^{Q100/+}$ | 8 | 10 | Saline | female | 4 | 133.0 | 4.1 |

**Supplementary Table 1: Cohort characteristics of *Htt* ASO treated *Htt*$^{Q111/+}$ and *Atxn2*$^{Q100/+}$ mice.** Importantly, inherited allele sizes did not differ significantly among treatments in the *Htt*$^{Q111/+}$ or *Atxn2*$^{Q100/+}$ cohorts (ANOVA treatment effect for *Htt*$^{Q111/+}$ cohorts: $F_{(2, 54)} = 0.42$ p = 0.66); and *Atxn2*$^{Q100/+}$ cohorts: $F_{(2,15)} = 1.69$, p =0.22). Inherited allele sizes were determined from tail tissue taken at weaning.

| Tissue | Age of IPTG withdrawal (months) | Age of mHTT suppression (months) | Sacrifice age (months) | N (female, male) | Average Inherited Allele Size (CAG no.) | Inherited Allele Size Standard Deviation |
|---|---|---|---|---|---|---|
| Liver | No IPTG | 0 - 6 | 6 | 7, 6 | 154.4 | 2.6 |
| Liver | 2 | 2 - 6 | 6 | 6, 7 | 155.8 | 4.1 |
| Liver | never | No suppression | 6 | 4, 5 | 156.4 | 3.8 |
| Liver | No IPTG | 0 - 12 | 12 | 8, 6 | 153.9 | 2.9 |
| Liver | 2 | 2 – 12 | 12 | 6, 7 | 155.3 | 2.5 |
| Liver | 8 | 8 – 12 | 12 | 7, 7 | 154.1 | 2.4 |
| Liver | Never | No suppression | 12 | 7, 7 | 155 | 2.5 |
| Cortex | No IPTG | 0 - 6 | 6 | 5, 5 | 154.7 | 2.7 |
| Cortex | 2 | 2 - 6 | 6 | 5, 5 | 155.4 | 3.2 |
| Cortex | never | No suppression | 6 | 5, 5 | 156.9 | 4 |
| Cortex | No IPTG | 0 - 12 | 12 | 5, 5 | 154.2 | 3.3 |
| Cortex | 2 | 2 – 12 | 12 | 5, 5 | 155.8 | 2.1 |
| Cortex | 8 | 8 – 12 | 12 | 5, 5 | 154.5 | 2.6 |
| Cortex | Never | No suppression | 12 | 5, 5 | 155.7 | 2.7 |
| Striatum | No IPTG | 0 - 6 | 6 | 5, 5 | 154.7 | 2.7 |
| Striatum | 2 | 2 - 6 | 6 | 5, 5 | 155.4 | 3.2 |
| Striatum | never | No suppression | 6 | 5, 5 | 156.9 | 4 |
| Striatum | No IPTG | 0 - 12 | 12 | 5, 5 | 154.2 | 3.3 |
| Striatum | 2 | 2 – 12 | 12 | 5, 5 | 155.8 | 2.1 |
| Striatum | 8 | 8 – 12 | 12 | 5, 5 | 154.4 | 2.6 |
| Striatum | Never | No suppression | 12 | 5, 5 | 155.7 | 2.7 |

**Supplementary Table 2: Cohort characteristics of the *Htt*$^{LacO\text{-}Q140/+}$ mice.** Inherited allele sizes did not differ significantly among the mice used to analyze liver somatic expansion (ANOVA treatment effect at 6 months: $F_{(2, 32)} = 1.01$, $p = 0.38$; and 12 months: $F_{(3, 51)} = 0.91$, $p = 0.44$). Within each terminal timepoint, the same mice were used to quantify striatal and cortical somatic expansion indices, with no significant differences in the inherited allele sizes of the mice (ANOVA treatment effect at 6 months: $F_{(2, 27)} = 1.15$, $p = 0.33$; and 12 months: ANOVA for treatment effect: $F_{(3, 36)} = 0.92$, $p = 0.44$). Inherited allele sizes were determined from tail tissue taken at weaning.
