## Supplementary Figures 1-3 for "Huntingtin lowering reduces somatic instability at CAG-expanded loci"

### Supplementary Material

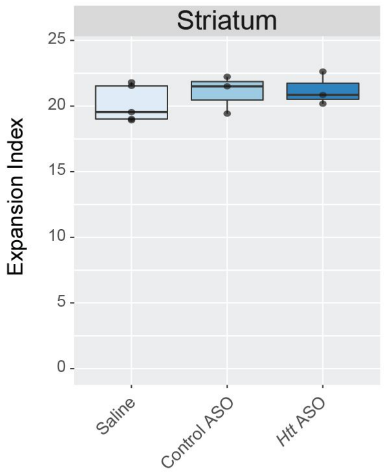


**Supplementary Figure 1: Peripheral HTT lowering does not impact expansion of the *Htt*$^{Q111}$ CAG tract in the striatum.** We have previously shown that peripheral *Htt* ASO treatment from 2 to 10 months of age does not reduce striatal HTT levels [16]. Unsurprisingly, striatal instability in those mice is unaffected by IP injection of *Htt* ASOs, which do not cross the blood brain barrier (ANOVA treatment effect: $F_{(2,8)} = 0.68$, $p = 0.53$). A 10% threshold was applied to calculate striatal expansion indices.

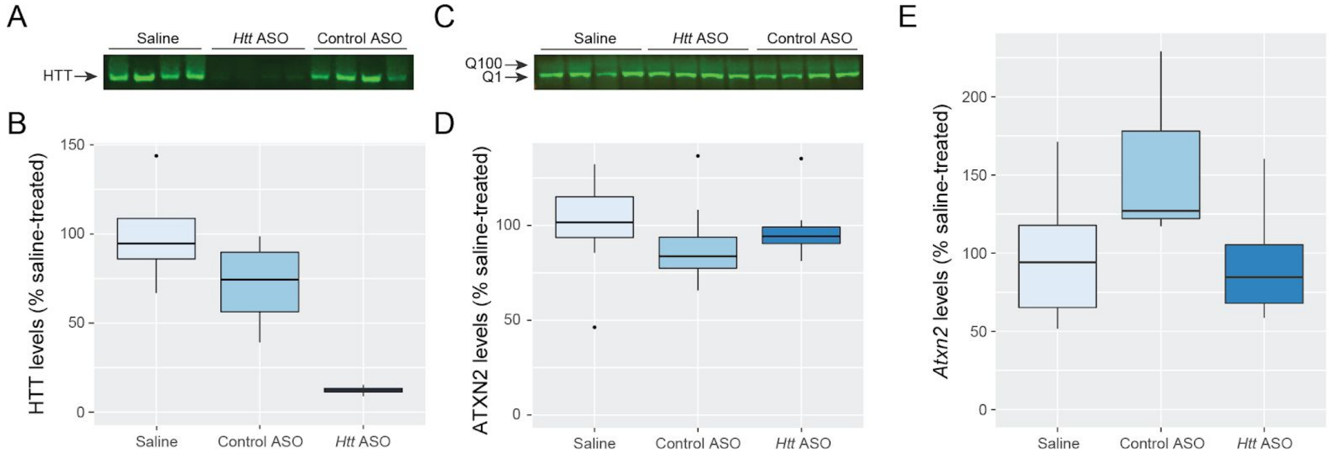


**Supplementary Figure 2: Hepatic HTT levels are reduced in *Atxn2*$^{Q100/+}$ mice by *Htt* ASO treatment, while ataxin-2 levels are unchanged.** A: Liver HTT levels were assessed via western blot after 8 months of treatment with *Htt* ASO, control ASO, or saline. B: *Htt* ASO treatment reduced HTT protein levels *Atxn2*$^{Q100/+}$ mouse livers (ANOVA for treatment effect: $F_{(2, 9)}$ = 13.8, p=1.8e-3). HTT protein signal was normalized to total protein signal on a per lane basis and expressed as a percent of the average HTT levels of saline-treated mice. C: Western blot showing mutant (Q100-ATXN2) and wild type (Q1-ATXN2) protein levels in the livers of 10-month-old *Atxn2*$^{Q100/+}$ mice following 8 months of ASO treatment. Consistent with previous reports [22], we observe reduced protein abundance of Q100-ATXN2 compared to Q1-ATXN2 in heterozygous mice. D: Quantification of total ATXN2 levels - western blot signal for each mouse was normalized to the total protein stain of the corresponding lane and expressed as a percent of the average total ATXN2 levels of saline-treated mice within each blot. Two western blots were run, bringing the total N per treatment arm to 8 mice. No differences in total ATXN2 (Q1- and Q100-ATXN2 species quantified together) were observed between treatment groups (ANOVA for treatment effect: $F_{(2, 21)}$ = 0.47, p = 0.63). E: Liver *Atxn2* levels were assessed via qRT-PCR, and did not differ between treatment groups (ANOVA for treatment effect: $F_{(2,10)}$ = 1.81, p = 0.21). *Atxn2* levels were normalized to *β-Act* on a per well basis and expressed as a percent saline-treated mice.

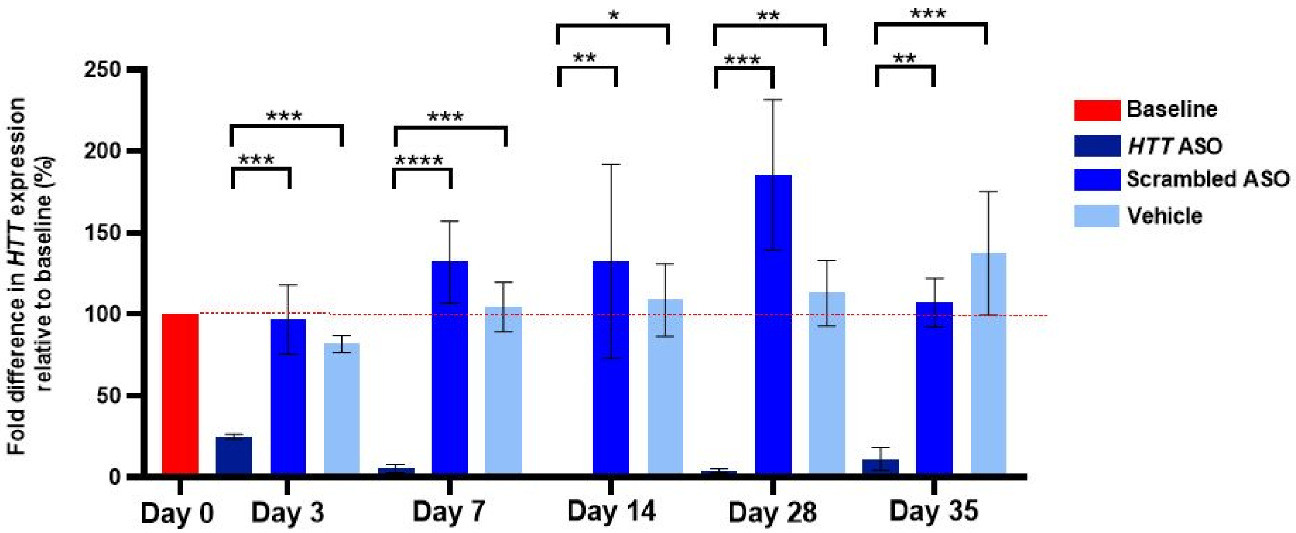


**Supplementary Figure 3: *HTT* levels in HD MSN-containing cultures are reduced by *HTT* ASO treatment.** *HTT* expression levels in HD MSN-containing cultures were quantified via qRT-PCR after 0, 3, 7, 14, 28 and 35 days of treatment with 40 µM *HTT* ASO, 40 µM scrambled ASO or vehicle. *HTT* expression levels were expressed as a percentage of the *HTT* levels in the untreated baseline sample, which is set as 100%. Treatment with *HTT* ASO reduces *HTT* expression compared to scrambled ASO and vehicle in HD MSN-containing cultures (ANOVA for treatment effect: $F_{(15,32)} = 16.24$, p = <0.0001). Data from each treatment group represents one biological replicate (one MSN differentiation) and three technical replicates. Error bars represent mean ± standard deviation. *: p= <0.05; **: p = <0.01; ***: p = <0.001; ****: p = <0.0001.
